# MpmiR319 promotes gemma/gemma cup formation in the liverwort *Marchantia polymorpha*

**DOI:** 10.1101/2023.09.29.560093

**Authors:** Kazutaka Futagami, Masayuki Tsuzuki, Yuichiro Watanabe

## Abstract

After microRNAs (miRNAs) were initially detected as approximately 21-nt sequences in plants, miRNA-mediated regulation of target gene expression was characterized in many cases. The sequences of miR159/319 family miRNAs are highly conserved and their development-related roles have been comprehensively characterized in dicot and monocot plants. However, their effects in basal land plants remain relatively unknown. We used a CRISPR/Cas9 system to edit genome sequences at Mp*MIR319a* and/or Mp*MIR319b* loci. Mutant *mir319b* lines developed relatively few gemma cups, while mutant *mir319a* lines produced relatively few gemmae in each gemma cup. Earlier 5′-RACE or degradome analyses suggested that Mp*miR319* targets Mp*RKD* and Mp*R2R3-MYB21* transcripts and suppresses expression. To determine whether these candidate genes influence the *mir319a/mir319b* phenotype we constructed miR319-resistant Mp*RKD* (mMp*RKD*) and Mp*R2R3-MYB21* (mMp*R2R3-MYB21*) by decreasing the complementarity to miR319. We also introduced the genes into wild-type liverwort and screened for phenotypic changes. We observed that mMp*RKD* resulted in gemma/gemma cup-less liverwort mutants, but mMp*R2R3-MYB21* did not. No major adverse effects were detected during male and female reproductive development induced by far-red illumination. These results indicate that miR319 influences gemma formation mainly by repressing Mp*RKD* expression rather than Mp*R2R3-MYB21* expression.

**Highlight:** This study demonstrates Mp*miR319* expression affects gemma cup and gemma formation during *Marchantia polymorpha* vegetative propagation by suppressing its target mRNAs.

## Introduction

The complementarity to their target mRNA sequences is generally greater for plant microRNAs (miRNAs) than for animal miRNAs (Rhoades et al., 2002)(Lin and Bowman, 2018). Each plant miRNA can target a limited number of mRNAs. Accordingly, mRNA(s) targeted by each miRNA can be predicted by searching genome sequence data for a sequence complementary to the miRNA sequence of interest. Recent studies revealed that land plants have 300–500 lineage-specific diverse miRNAs (Axtell et al., 2007)(Lin and Bowman, 2018)(Pietrykowska et al., 2022). Earlier *in vivo* studies on miRNAs conserved among land plants showed that their target mRNAs are transcribed from highly conserved cognate genes across phyla (Axtell et al., 2007)(Tsuzuki et al., 2019)(Alaba et al., 2015)(Pietrykowska et al., 2022). Such conserved miRNA–target mRNA modules may form a core set of functionally and biologically important gene regulatory circuits in land plants. Moreover, as few as seven miRNA families are highly conserved from the liverwort *Marchantia polymorpha* and the moss *Physcomitrella patens* to angiosperms (Axtell et al., 2007)(Tsuzuki et al., 2016)(Lin et al., 2016b)(Pietrykowska et al., 2022). However, most liverwort miRNAs have not been studied sufficiently to determine how common or specific they are among land plants. Additional experimental data are needed to substantiate the biological significance of highly conserved miRNAs in basal land plants. Among conserved miRNAs, miR159/319 family members are unique because their target mRNA species may have changed during phylogenetic diversification (Palatnik et al., 2007). In Arabidopsis, miR319 preferentially targets mRNAs of *TEOSINTE BRANCHED1*, *CYCLOIDEA*, and *PROLIFERATING CELL FACTOR* genes (Martín-Trillo and Cubas, 2010)(Kosugi and Ohashi, 1997). These genes encode basic-helix-loop-helix transcription factors that redundantly affect cotyledon boundary specification and leaf serration (Palatnik et al., 2003)(Koyama et al., 2017).

*Marchantia polymorpha* is a liverwort that is a valuable emerging model species used in studies on land plant diversification, which have generated increasing amounts of information and resources for comparative evolution research (Bowman et al., 2017). In *M. polymorpha*, Mp*miR319* targets mRNAs of *MpR2R3-MYB21* (Arabidopsis MYB33-like transcription factor belonging to the GAMYB family) (Lin et al., 2016b)(Tsuzuki et al., 2016) and RWP-RK DOMAIN (RKD)-containing transcription factor genes (Lin et al., 2016b) according to a 5′-RACE or degradome analysis. Thus, Mp*miR319* may have some regulatory effects on such candidate target genes, but this possibility remains to be experimentally verified.

## Materials and methods

### Plant materials and growth conditions

Takaragaike-1 (Tak-1, male) and Takaragaike-2 (Tak-2, female) were used as wild-type *M. polymorpha* samples (Ishizaki et al., 2008). Plants were grown asexually on half-strength Gamborg’s B5 medium (FUJIFILM Wako Pure Chemical Co., Osaka, Japan) (Gamborg et al., 1968) containing 0.5 g/l 2-morpholinoethanesulfonic acid (MES, Tokyo Chemical industry Co., Tokyo) and 1% (w/v) agar (STAR Agar L-grade, RIKAKEN, Tokyo) under continuous white fluorescent light at 22 °C. To induce the sexual reproductive phase, plants were grown at 22 °C under continuous light conditions provided by a mixture of white fluorescent light and far-red (FR) light-emitting diodes (OSXXXXT3C1E, OptoSupply, Hong Kong; peak emission at 730 nm, with a full width at half-maximum of 20 nm).

### Transformation and mutant selection

Regenerating thalli (for *mir319a* or *mir319b* mutants) (Kubota et al., 2013) and a G-AgarTrap method (to generate gMp*RKD*, mMp*RKD*, gMp*MYB21*, and mMp*MYB21* lines) (Tsuboyama and Kodama, 2018) were used to transform *M. polymorpha*. Briefly, transformed plants were selected on half-strength Gamborg’s B5 medium containing 0.5 g/l MES, 1% (w/v) agar, 100 µg/ml cefotaxime and 10 µg/ml hygromycin (for pMpGWB101 and pMpGE010), or 100 µg/ml cefotaxime and 0.5 µM chlorsulfuron (for pMpGWB301 and pMpGE011). After screening T_1_ plants on drug-supplemented media, we obtained approximately 10 candidate lines for each guide RNA (gRNA) in the male and female genetic backgrounds.

### Genomic DNA extraction

Genomic DNA was extracted from Tak-1 thalli using the DNeasy Plant Mini kit (Qiagen, Hilden, Germany) according to the manufacturer’s recommendations.

### Plasmid construction for the CRISPR/Cas9 system

We employed the CasFinder Tool (https://marchantia.info/tools/casfinder/) to design gRNAs to target Mp*miR319a* and/or Mp*miR319b* loci. The pMpGE_En03 vector (Sugano et al., 2018) was used to clone oligonucleotide pairs to express gRNAs: pMpGE_En03-gRNA1, pMpGE_En03-gRNA3, and pMpGE_En03-gRNA4. pMpGE010 and pMpGE011 were used for Cas9 expression (Sugano et al., 2018).

The *mir319a^ge^*, *mir319b^ge^*, and *mir319ab^ge^* mutants were selected as described previously (Tsuzuki et al., 2019). Using genomic DNA as the template, *MIR319A* and *MIR319B* sequences were amplified by PCR using PrimeSTAR HS DNA polymerase (TaKaRa Bio, Kusatsu, Japan) and the primer pairs MpMIR319aF1 and MpMIR319a-R1 or MpMIR319bF2 and MpMIR319b-R2. Amplified products were sequenced according to the Sanger method. Details regarding the PCR and genomic sequencing primers are listed in Table S1. Mp*MIR319A* and Mp*MIR319B* genomic sequences in *mir319a^ge^*, *mir319b^ge^*, and *mir319ab^ge^* mutant plants (T_1_ generation) are provided in Figure 1.

**Fig. 1.**
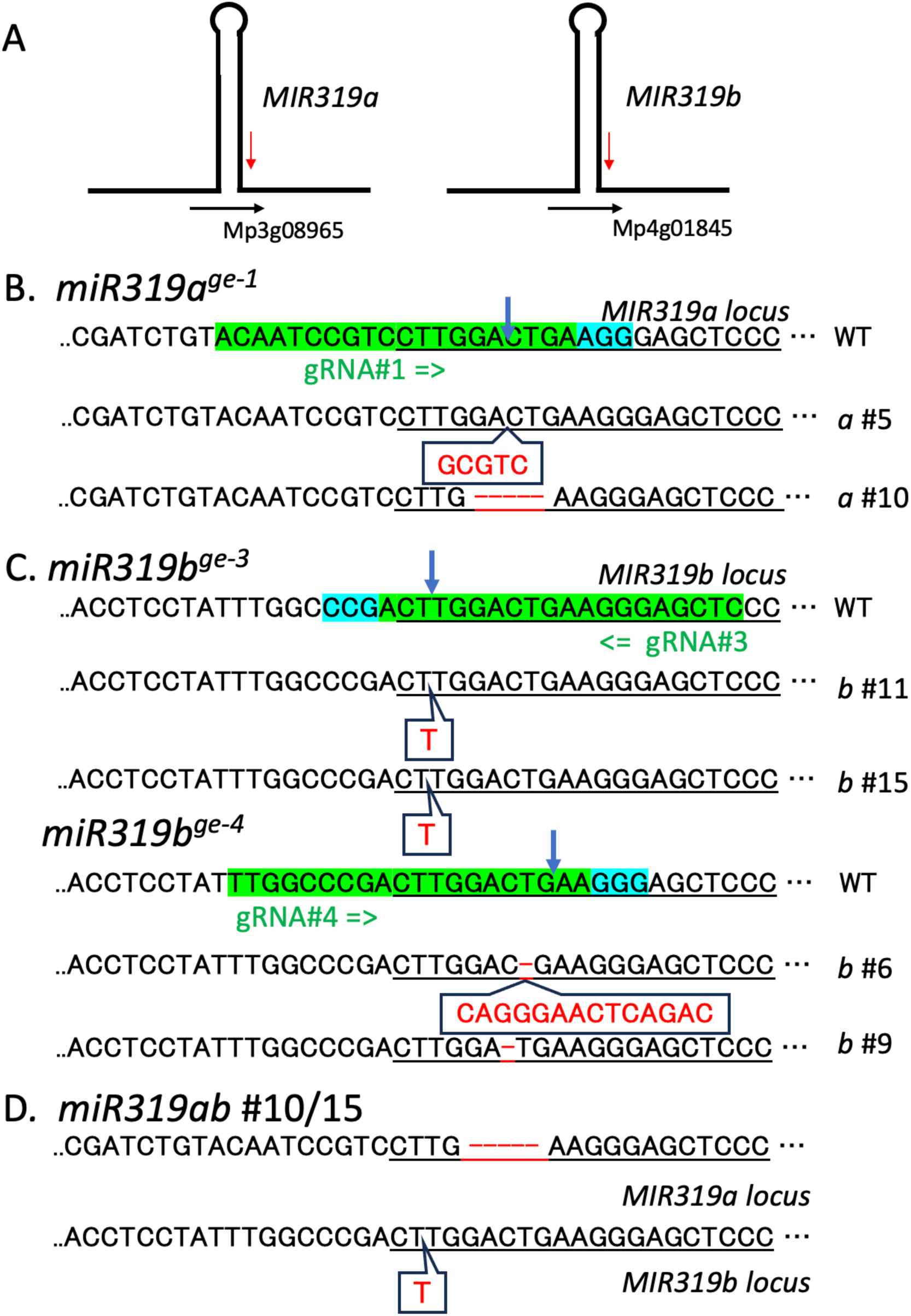
(A) Primary *MIR319a* or *b* RNAs are predicted to be transcribed from two annotated loci in the *M. polymorpha* genome. (B) The wild-type (WT) *MIR319a* locus sequence targeted by gRNA#1 is presented. Mutants #5 and #10 were analyzed further. (C) The WT *MIR319b* locus sequence targeted by gRNA#3 or #4 is presented. Mutants #11 and #15 from the gRNA#3-treated pool and #6 and #9 from the gRNA#4-treated pool were analyzed further. In the superscript *ge-x*, *x* indicates the gRNA species used to create the mutant. (D) Changes in the *MIR319a* and *b* genomic loci in the double knockout line *miR319ab #10/15*. The *MIR319a* locus had a 5-bp deletion and the *MIR319b* locus had a 1-bp insertion, similar to *miR319a^ge-1^* #10 and *miR319b^ge-3^*. For simplicity, we named the line *miR319ab #10/15*.

### Construction of miR319-resistant Mp*RKD* and Mp*R2R3-MYB21* transgenes

To obtain the wild-type *RKD* gene segment, the genomic sequence from the Mp*RKD* promoter region (3,827 bp upstream of the translation initiation codon) to the 3′ untranslated region (UTR) (568 bp downstream of the translation termination codon) was amplified by PCR using Tak-1 genomic DNA, PrimeSTAR HS DNA polymerase (TaKaRa Bio), and primers IF-Hind-pRKD-F and Inf-gMpRKD-Sal-R. To create the miR319-resistant Mp*RKD* gene segment, two PCR fragments were amplified using Tak-1 genomic DNA as well as the IF-Hind-pRKD-F and mRWP_RK-R primer set and the mRWP_RK-F and Inf-gMpRKD-Sal-R primer set to specifically mutate the miR319 target sequence in the Mp*RKD* 3′ UTR sequence. The In-Fusion Cloning kit (TaKaRa Bio) was used to insert the PCR fragments into the pMpGWB300 vector that had been cleaved by *Hin*dIII and *Sal*I to generate the pMpGWB300:proMpRKD:gMpRKD and pMpGWB300:proMpRKD:mMpRKD recombinant plasmids.

To obtain the wild-type *MYB21* gene segment, the Mp*MYB21* promoter region (3008 bp upstream of the translation initiation codon) to the initiation codon, was first amplified by PCR using Tak-1 genomic DNA, PrimeSTAR HS DNA polymerase, and primers Hind-pMpMYB21-3000-F and Sal-pMpMYB21-R, cloned into pMpGWB100 after both being restricted with *Hin*dIII and *Sal*I to obtain pMpGWB100-pMYB21. Wild-type *MYB21* gene segment (gMYB21) from halfway of promoter region to the termination codon (4244 bp) were amplified using Tak-1 genomic DNA and primers pMYB33-459F and Bam-MpMYB33-R2. To create the miR319-resistant Mp*MYB21* gene segment, two PCR fragments were amplified using gMYB21 fragment as well as the pMYB33-459F and mMYB33-R set and the mMYB33-F and Bam-MpMYB33-R2 primer set to specifically mutate the miR319 target sequence in the Mp*MYB21* coding sequence. The obtained PCR fragments were mixed and subjected to the next PCR reaction using primers pMYB33-459F and Bam-MpMYB33-R2 to obtain miR319 resistant MYB21 gene body fragment (mMYB21). The gMYB21 and mMYB21 fragments were cloned separately into the pMpGWB100 vector, after being compatibly restricted with *Nru*I and *Bam*HI, to generate the pMpGWB100:pgMp*MYB21* and pMpGWB100:pmMp*MYB21*.

*M. polymorpha* was transformed with the recombinant plasmids as described previously (Tsuboyama et al., 2018a)(Tsuboyama and Kodama, 2018).

## Results

### Guide RNA-mediated editing of *MIR319* genes

A previous study revealed *M. polymorpha* has a relatively simple genome because of limited gene redundancy (Bowman et al., 2017). *M. polymorpha* has two *MIR319* coding loci, both of which produce an identical mature miRNA sequence (5′-_p_CUUGGACUGAAGGGAGCUCCC_OH_-3′) (Lin et al., 2016b; Tsuzuki et al., 2016). The two *MIR319* loci have been annotated (*M. polymorpha* genome ver.6.1) as Mp*MIR319A* (Mp3g08965) and Mp*MIR319B* (Mp4g01845) (Fig. 1A).

We focused on whether Mp*MIR319a* and/or Mp*MIR319b* loci are functional. To analyze the possible biological functions of miR319 in *M. polymorpha*, we edited two annotated Mp*miR319* coding sequences to obtain mutants, which were examined for phenotypic changes. Specifically, we designed gRNA#1 for *MIR319A* (Fig. 1B) and gRNA#3 and #4 for *MIR319B* (Fig. 1C) and constructed vectors to introduce indel mutations at the respective *MIR319* loci using a CRISPR/Cas9 genome-editing system (Sugano et al., 2014)(Sugano et al., 2018). We adopted an *Agrobacterium*-mediated transformation protocol to introduce the vectors into Tak-1 thalli to express the gRNAs and Cas9 (Sugano et al., 2014)(Tsuboyama et al., 2018b). After the drug-based selection of G_1_ plants, we amplified the targeted genomic sequences from selected G_1_ thalli and analyzed the PCR products via Sanger sequencing to confirm indel mutations were present at the target loci. Multiple lines with indel mutations at the Mp*MIR319a* or Mp*MIR319b* locus were generated. Several representative lines with similar phenotypes were selected (Fig. 1B and C) for the subsequent analysis. The selected mutants with indel mutations at the Mp*mir319a* locus were designated as *mir319a^ge-1^ a* #5 and *mir319a^ge-1^ a* #10, whereas the selected mutants with indel mutations at the Mp*mir319b* locus were designated as *mir319b*^ge-3^ *b* #11, *mir319b*^ge-3^ *b* #15, *mir319b*^ge-4^ *b* #6, and *mir319b*^ge-4^ *b* #9.

### *miR319a* and *miR319b* mutant thalli exhibited abnormal gemma cup production

Although the *mir319a^ge^* and *mir319b*^ge^ mutant plants grew as fast as the wild-type Tak-1 plants, they produced gemma cups less frequently than Tak-1 plants (Fig. 2). More specifically, compared with wild-type Tak-1 plants, significantly fewer *mir319a^ge-1^* (*a* #5 and *a* #10) and *mir319b*^ge-3^ (*b* #11 and *b* #15) plants produced gemma cups (approximately half as many). Gemma cup production was most inhibited in the *mir319b*^ge-4^ (*b* #6 and *b* #9) plants. We speculated that both *mir319a* and *miR319b* alleles somehow affect gemma cup formation during dorsal organ formation in thalli. In addition, *mir319a^ge-1^* (*a* #5 and *a*#10) mutants produced fewer gemmae in each gemma cup than the wild-type plants (Fig. 3), whereas the *mir319b*^ge-3^ (*b* #11 and *b* #15) mutants and the wild-type plants did not differ in terms of the number of gemmae in each gemma cup (Fig. 3). These observations suggest that miR319 contributes to the formation of gemmae and gemma cups in *M. polymorpha* by regulating target gene expression.

**Fig. 2.**
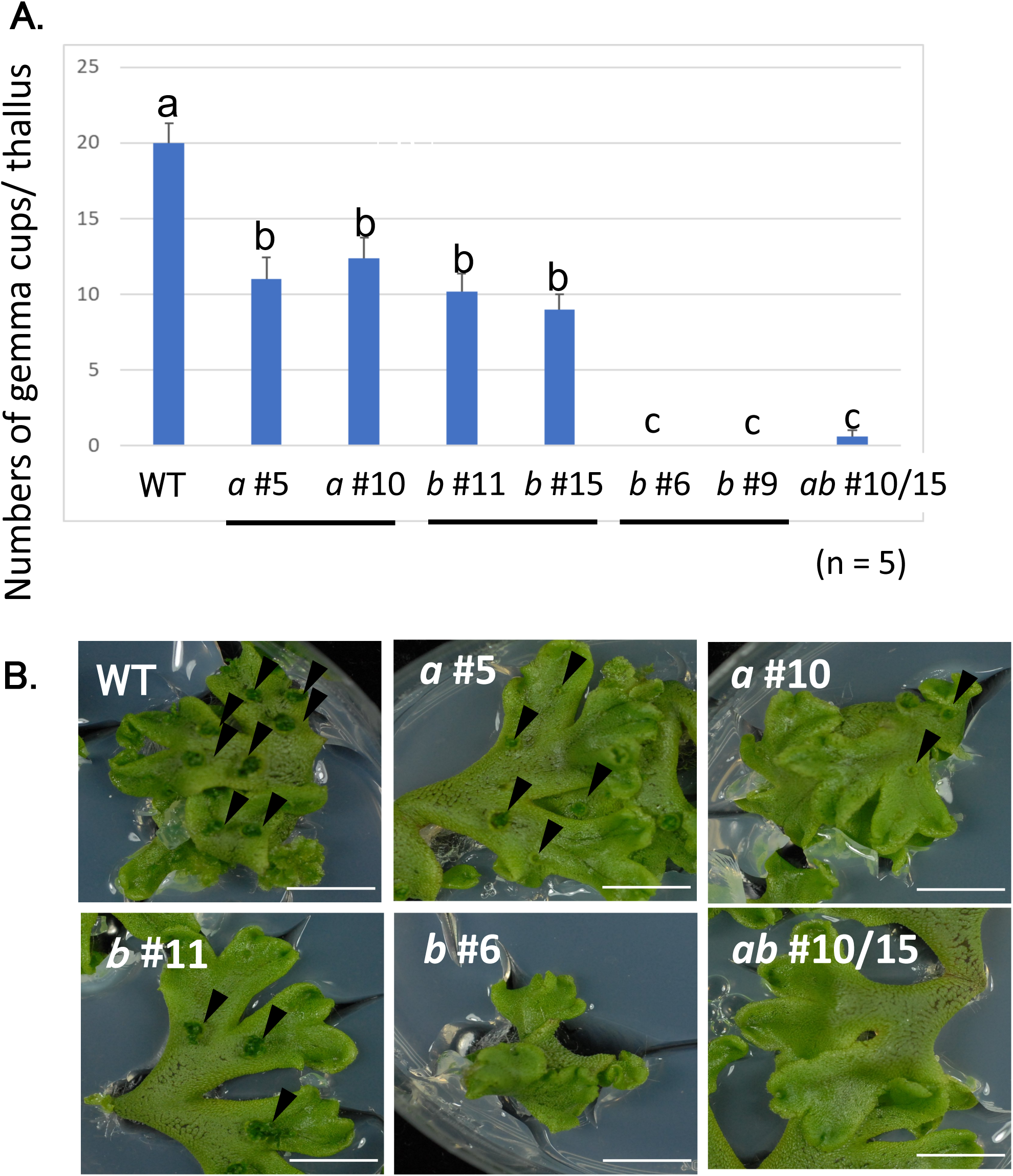
(A) Disruption of the Mp*MIR319a* and/or *b* loci decreased the number of gemma cups. Five plants were analyzed and the mean number of gemma cups per thallus is presented. Tukey–Kramer’s honestly significant difference test was performed to analyze the variance. Statistically significant differences are indicated by different letters (a, b, c; *P* < 0.05). (B) Appearance of different thalli. Bar = 1 cm. WT: Tak-1 wild-type plant, #5: *miR319a^ge-1^* #5, #10: *miR319a^ge-1^* #10, #11: *miR319b^ge-3^* #11, #15: *miR319b^ge-3^* #15, #6: *miR319b^ge-4^* #6, #10/15: *miR319ab #*10/15.

**Fig. 3.**
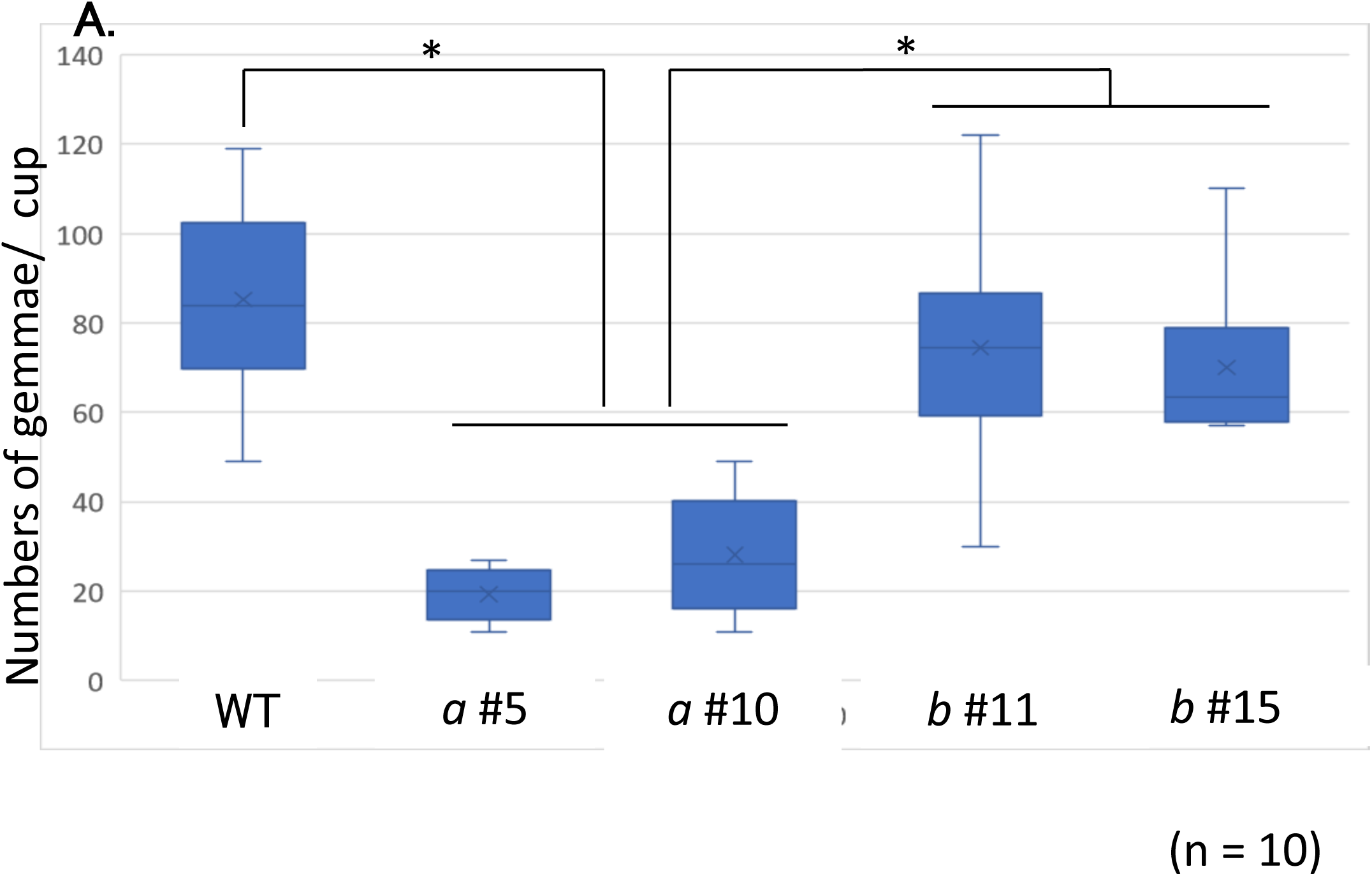
Disruption of the Mp*MIR319a* and/or *b* loci inhibited gemma formation. (A) Number of gemmae formed in each gemma cup. \**P* < 0.05 (Tukey method). (B) Appearance of a gemma cup. Bar = 1 cm. WT: Tak-1 wild-type plant, #5: *miR319a^ge-1^* #5, #10: *miR319a^ge-1^* #10, #11: *miR319b^ge-3^* #11, #15: *miR319b^ge-3^* #15.

We also created similar *mir319a^ge^* and *mir319b*^ge^ mutants using female Tak-2 plants. A decrease in gemma cup formation was also detected in the Tak-2-based mutants (data not shown). Thus, we focused mainly on lines with a Tak-1 genetic background to study the miR319 function during vegetative growth stages.

### Knockout of both *miR319a* and *miR319b* loci did not result in additive defects

The *mir319a* and *mir319b* knockout lines exhibited similar phenotypic defects in terms of the production of gemma cups/gemmae. We analyzed whether the two loci additively affect liverwort vegetative organ formation by creating and characterizing double knockout mutants. We were unable to obtain a double mutant via a single transformation involving two vectors. Thus, we targeted the *MIR319a* locus of *mir319b^ge-3^ b* #15 with pMpGE_En03-gRNA1. After several trials, we obtained a double knockout line that had a 5-base deletion in the *MIR319a* locus identical to *mir319a^ge-1^ a* #10 in addition to a 1-base addition in the *MIR319b* locus (Fig. 1D). For brevity, the mutant line was named *miR319ab #10/15* (Fig. 1D). In terms of the number of developed gemma cups and gemmae, the *miR319ab #10/15* mutant phenotype did not differ from the *miR319a* and *mir319b* mutant phenotypes (Fig. 2, *ab* #10/15).

### Regulation of Mp*RKD* by miR319 influences gemma cup formation

An earlier degradome analysis indicated that Mp*RKD* (formerly *RWP-RK DOMAIN-CONTAINING 1*) and Mp*MYB33* mRNAs are targeted by Mp*miR319* (Lin et al., 2016a). In the current study, we assessed whether the susceptibility of either or both of these mRNA genes to miR319 affects gemma/gemma cup formation. Mp*RKD* encodes an RWP-RK domain-containing transcription factor that is an evolutionarily conserved regulator of germ cell differentiation (Koi et al., 2016)(Rövekamp et al., 2016). Additionally, it was reported that Mp*RKD* is expressed on gemma cup rims (Koi et al., 2016) and is necessary for normal gemma cup growth (Rövekamp et al., 2016). Accordingly, we examined whether downregulated Mp*RKD* expression due to miR319 is linked to gemma/gemma cup formation. We constructed a mutated *RKD* coding sequence that is unaffected by miR319.

The 21-nt sequence complementary to miR319 started from 4 nt downstream of the stop codon in the *M. polymorpha RKD* (Mp*RKD*) mRNA sequence. We constructed a miR319-resistant form of the Mp*RKD* gene (designated as mMp*RKD*; Fig. 4A) by introducing a 12-base substitution to decrease the complementarity to Mp*miR319*. The mMp*RKD* gene was placed downstream of its own 3.8 kb promoter sequence in the pMpGWB300 vector (Ishizaki et al., 2015). As a control, we cloned wild-type Mp*RKD* (designated as gMp*RKD*) in the same vector. The mMp*RKD* gene produces *RKD* mRNA that is insensitive to miR319, whereas gMp*RKD* mRNA is sensitive to miR319. The gMp*RKD* and mMp*RKD* genes were introduced into separate Tak-1 wild-type plants to obtain gMp*RKD* and mMp*RKD* lines. Both gMp*RKD* and mMp*RKD* plants had similar thallus growth rates, but the mMp*RKD* plants failed to produce gemma cups (Fig. 5: mMpRKD). The gemma cup-less phenotypes were similar to those of the *mir319a^ge-1^* or *miR319ab #10/15* mutants (Fig. 5; mir319ab #10/15). In contrast, the gMp*RKD* lines produced gemma cups similar to the wild-type Tak-1 control (Fig. 5; gMpRKD). Therefore, the sequences complementary to Mp*miR319* in Mp*RKD* mRNA are critical for gemma/gemma cup formation. Moreover, Mp*RKD* translation is likely suppressed by Mp*miR319*.

**Fig. 4.**
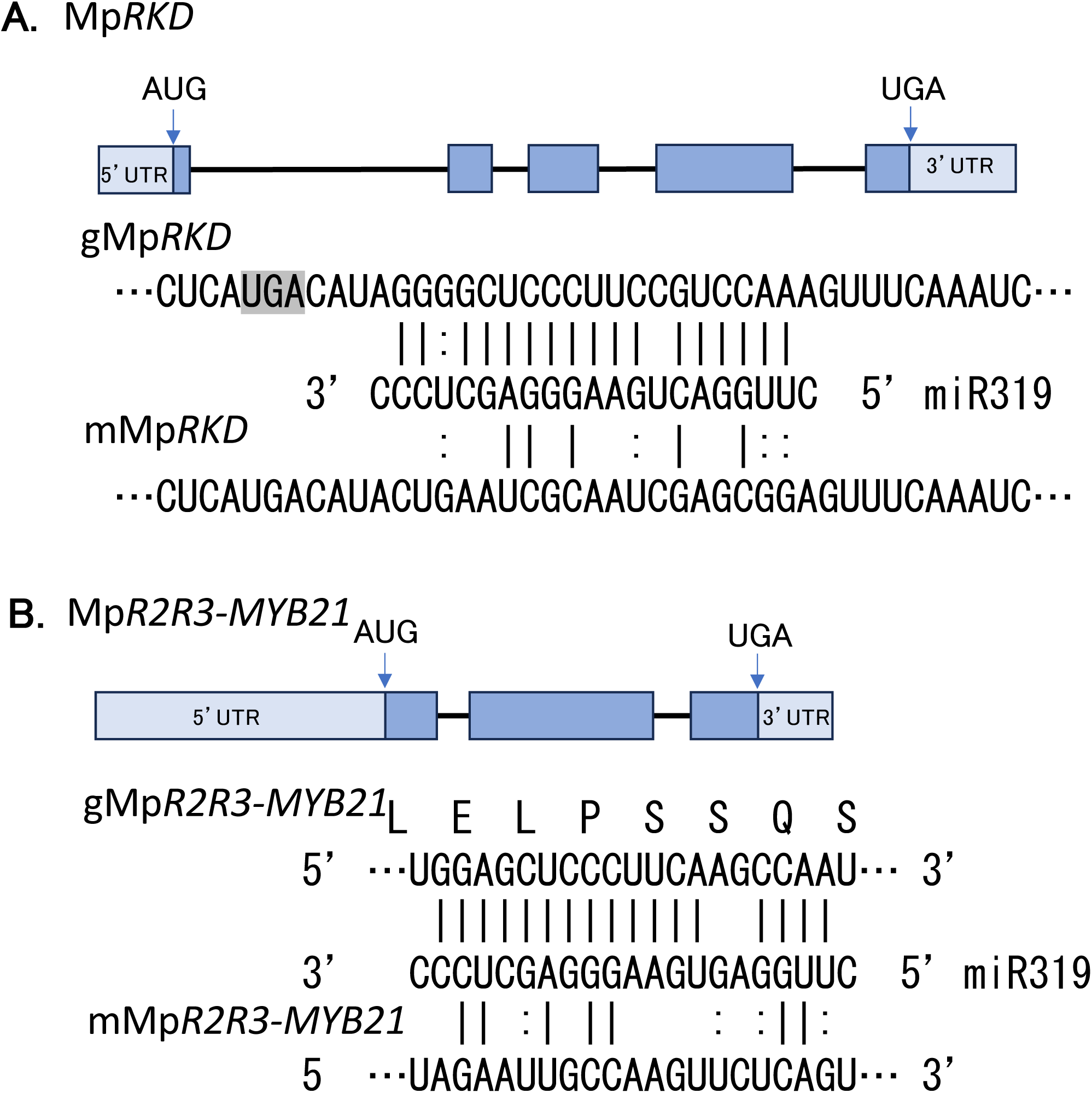
Exon–intron structures of Mp*RKD* and Mp*R2R3-MYB21* genes in the wild-type form (or sensitive form, with prefix g) and miR319-resistant form (with prefix m) of (A) Mp*RKD* and (B) mMp*R2R3-MYB21* genes. Vertical bars indicate conventional base-pairing, whereas double dots indicate G–U pairs. AUG and UGA are initiation and termination codons, respectively, of open reading frames. 5′UTR and 3′UTR refer to untranslated regions.

**Fig. 5.**
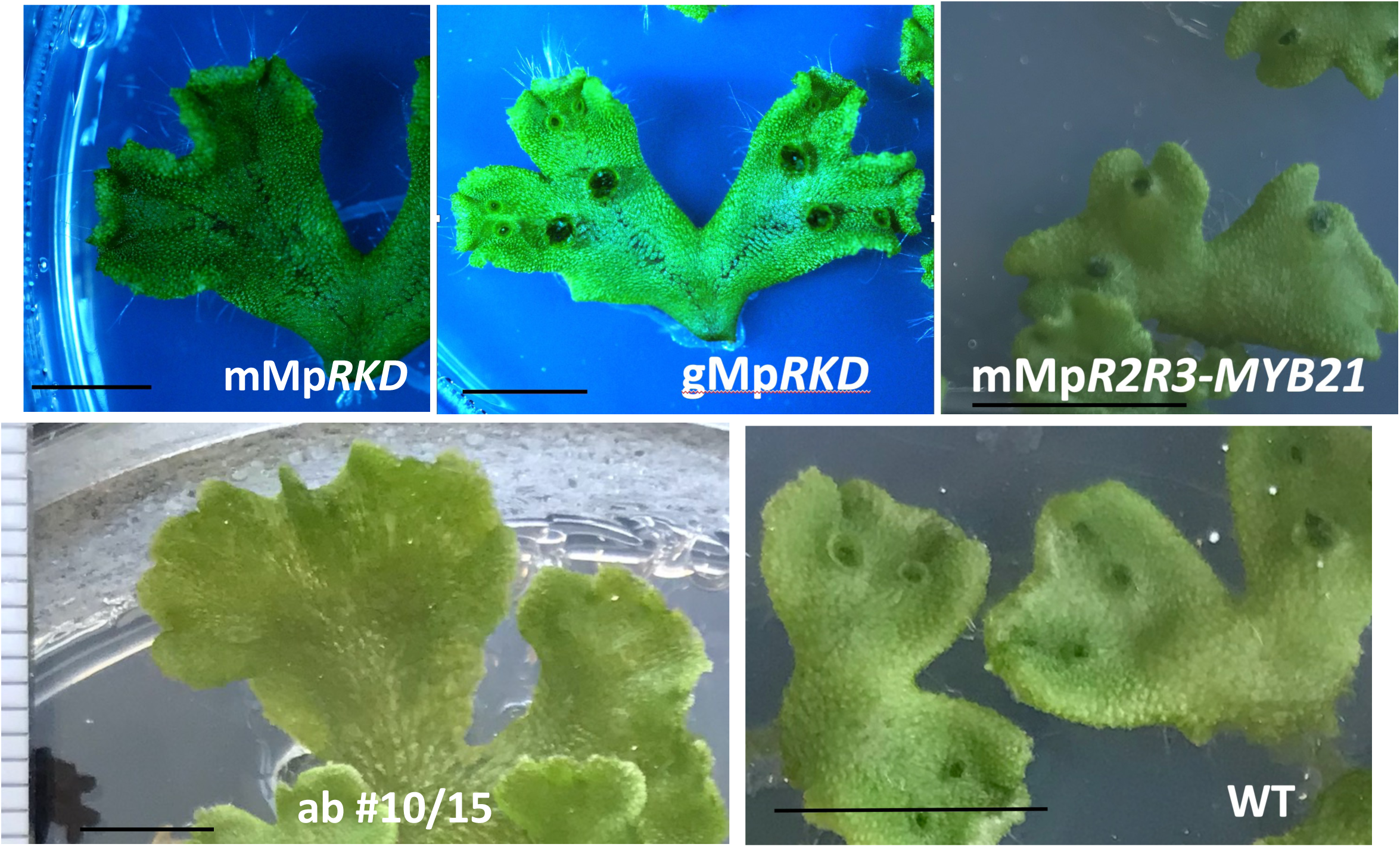
Introduction of the miR319-resistant mMp*RKD* gene suppressed gemma cup formation in wild-type thalli, but the introduction of gMp*RKD* and mMp*MYB21* genes did not.

As mentioned earlier, the miR319 target sequence is located in the 3′ UTR downstream of the *RKD* termination codon sequence. Thus, we designed another miR319-insensitive *RKD* gene (Mp*RKD*-Δ3′UTR) by removing the complementary sequences from the 3′ UTR without disturbing *RKD* translation (Fig. S1A). When we introduced Mp*RKD*- Δ3′UTR into liverwort, thalli were able to grow, but only a few gemma cups developed, similar to the mMp*RKD* liverwort (Fig. S1B).

### Gemma cup/gemma formation was unaffected by miR319-resistant Mp*MYB21*

According to a 5′-RACE or degradome analysis, Mp*MYB21* is also a potential target gene of miR319 (Tsuzuki et al., 2016)(Lin et al., 2016b). To test whether the regulation of *MYB21* expression by miR319 affects gemma cup formation, we created a miR319-resistant Mp*MYB21* (mMp*R2R3-MYB21* or m*MYB21*) that was placed under the control of its own promoter similar to the Mp*RKD* gene (Fig. 4B). mMp*R2R3-MYB21* was introduced into wild-type Tak-1 liverwort. The obtained m*MYB21* liverwort lines produced thalli with gemma cups/gemmae, but the thalli had shapes slightly different from the wild-type thalli (Fig. 5; mMpR2R3-MYB21). These results indicate that Mp*R2R3-MYB21* affects gemma cup/gemma formation substantially less than Mp*RKD* (if at all).

### mMp*RKD* plants developed deformed antheridiophores/archegoniophores, but normal sperm and eggs

It was previously reported that Mp*RKD*, which is expressed in developing eggs and sperm precursors, regulates egg and sperm formation in *M. polymorpha* (Koi et al., 2016)(Rövekamp et al., 2016). Thus, we analyzed germ cell, gametangia, and gametangiophore differentiation (Yamaoka et al., 2021)(Shimamura, 2016) in gMp*RKD* and mMp*RKD* liverworts. In response to FR illumination, the mMp*RKD* line with the Tak-1 genetic background developed male gametophytes (Fig. 6C) at nearly the same time as the gMp*RKD* liverwort (Fig. 6A). In addition, mMp*RKD* antheridiophores had almost normal stalked receptables, except for their irregular asymmetrical shape (Fig. 6C, D). Antheridia formed and protruded outward, but they were not surrounded by antheridiophores in mMp*RKD* lines (Fig. 6E, F). Sperm mobility was similar in the mMp*RKD*, wild-type Tak-1, and gMp*RKD* liverworts.

**Fig. 6.**
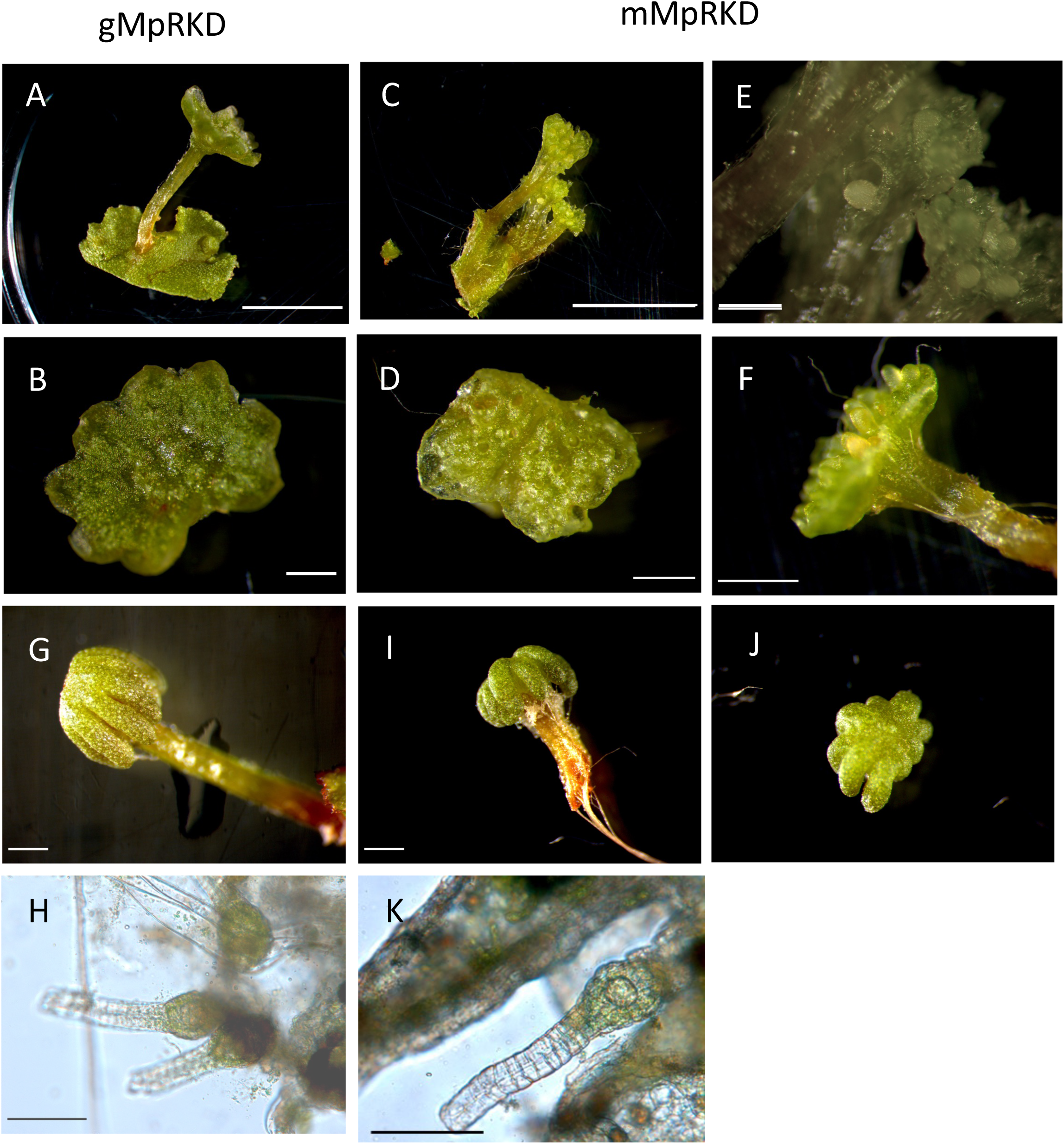
Introduction of the wild-type *RKD* gene (gMp*RKD*) induced normal gametophyte development in Tak-1 (A, B) and Tak-2 (G, H). Introduction of the miR319-resistant mutant *RKD* gene (mMp*RKD*) into Tak-1 induced the production of antheridiophores with stalked receptables with irregular asymmetrical shapes (C, D) and antheridiophores with exposed antheridia (E, F). Introduction of mMp*RKD* into Tak-2 induced the production of archegoniophores with short digits (I,J), but normal archegonia (K).

We also introduced gMp*RKD* and mMp*RKD* genes into wild-type Tak-2 liverwort and then checked whether archegonia/archegoniophore development was affected. We generated transgenic lines using two constructs (Fig. 4A). Under FR light, the mMp*RKD* liverwort in the Tak-2 genetic background developed female gametophytes as early as the gMp*RKD* liverwort, but its archegoniophores (Fig. 6I) differed from those of gMp*RKD* (Fig. 6G) or Tak-2 plants in terms of shape. mMp*RKD* archegoniophores had short digits (Fig. 6I,J), but produced normal archegonia. Furthermore, in terms of their appearance, the mMp*RKD* archegonia (Fig. 6K) were almost identical to the archegonia of gMp*RKD* (Fig. 6H) or wild-type Tak-2 liverwort.

## Discussion

When we profiled *M. polymorpha* miRNAs by small RNA-seq analysis, we tried to overexpress miR319 precursor RNAs in wild-type liverwort to clarify their biological functions. The ectopic expression of a miR319 precursor in liverwort adversely altered the development of gemma cups (Tsuzuki et al., 2016). We speculated that the balanced expression of miR319/target mRNAs is necessary for normal gemma cup development in *M. polymorpha*. Our RACE assay detected the cleavage of Mp*MYB21* mRNA, which encodes a homolog of Arabidopsis *MYB33*. An earlier RNA degradome analysis revealed Mp*RKD* (formerly *RWP-RK DOMAIN-CONTAINING 1*) mRNA is also cleaved by MpmiR319 (Lin et al., 2016b). Mp*RKD* was previously investigated using *Marchantia* mutants (Koi et al., 2016)(Rövekamp et al., 2016). These studies showed that Mp*RKD* is essential for normal germ cell differentiation in both male and female gametophytes, but it is also involved in the differentiation of non-reproductive cells surrounding the antheridia, developing egg, and sperm precursors (Koi et al., 2016)(Rövekamp et al., 2016). Mp*RKD* also influences gemma cup formation (Rövekamp et al., 2016). This gene is transcribed at the edge of gemma cups and the meristematic regions of apical notches (Koi et al., 2016)(Rövekamp et al., 2016). It remains unknown whether miR319 organizes gene regulatory networks with Mp*RKD* and further research will clarify.

The gemma cup formation has been histologically characterized. Founder gemmae arise directly from above the center of the apical notch of the thallus, which is followed by the formation of the cup rims surrounding gemmae (Shimamura, 2016). Our findings combined with the available data suggest that Mp*RKD* is involved in various cell proliferation stages, resulting in the formation of a variety of *M. polymorpha* tissues, including gemmae/gemma cups. Nevertheless, additional research is needed to clarify how gemma cups are organized by miR319-mediated gene regulatory networks.

There are several reports of mutants with defective pathways mediating gemma cup formation in thalli, including the RopGEF KARAPPO (Hiwatashi et al., 2019), GCAM1 (Yasui et al., 2019), and MpKAI2–ligand signaling (Komatsu et al., 2023) pathways. However, the pathway facilitating the formation of gemma cups from the apical notch has not been fully determined. Solly *et al*. showed that growth rate heterogeneity determines the flattened shape of *Marchantia* thalli and that the basic shapes are due to growth rate differences specified by apical notch activities (Solly et al., 2017). Recently, Romani *et al*. comprehensively revealed the activities of approximately 450 putative transcription factor gene promoters by analyzing the respective reporter fusion constructs (Romani et al., 2024). They generated a map of expression domains for each transcription factor gene related to gemma growth and identified more than 100 promoters that are active specifically in the apical notch. The mechanism underlying the formation of gemma cups may be much more complex than expected.

The *mir319a* and *mir319b* loci differentially contribute to gemma cup formation, possibly because of some differences in cell-specific transcription or processing efficiency, but this possibility will need to be tested in future studies. In the moss *P. patens*, miR159/319, which are encoded in the genome, may target mRNAs for cyclin domain proteins and two MYB transcription factors (Axtell et al., 2007). Moreover, miR159/319 families are highly conserved from bryophytes to angiosperms, but their biological functions might be more diverse than we initially imagined.

## Supplementary Data

**Table S1.**
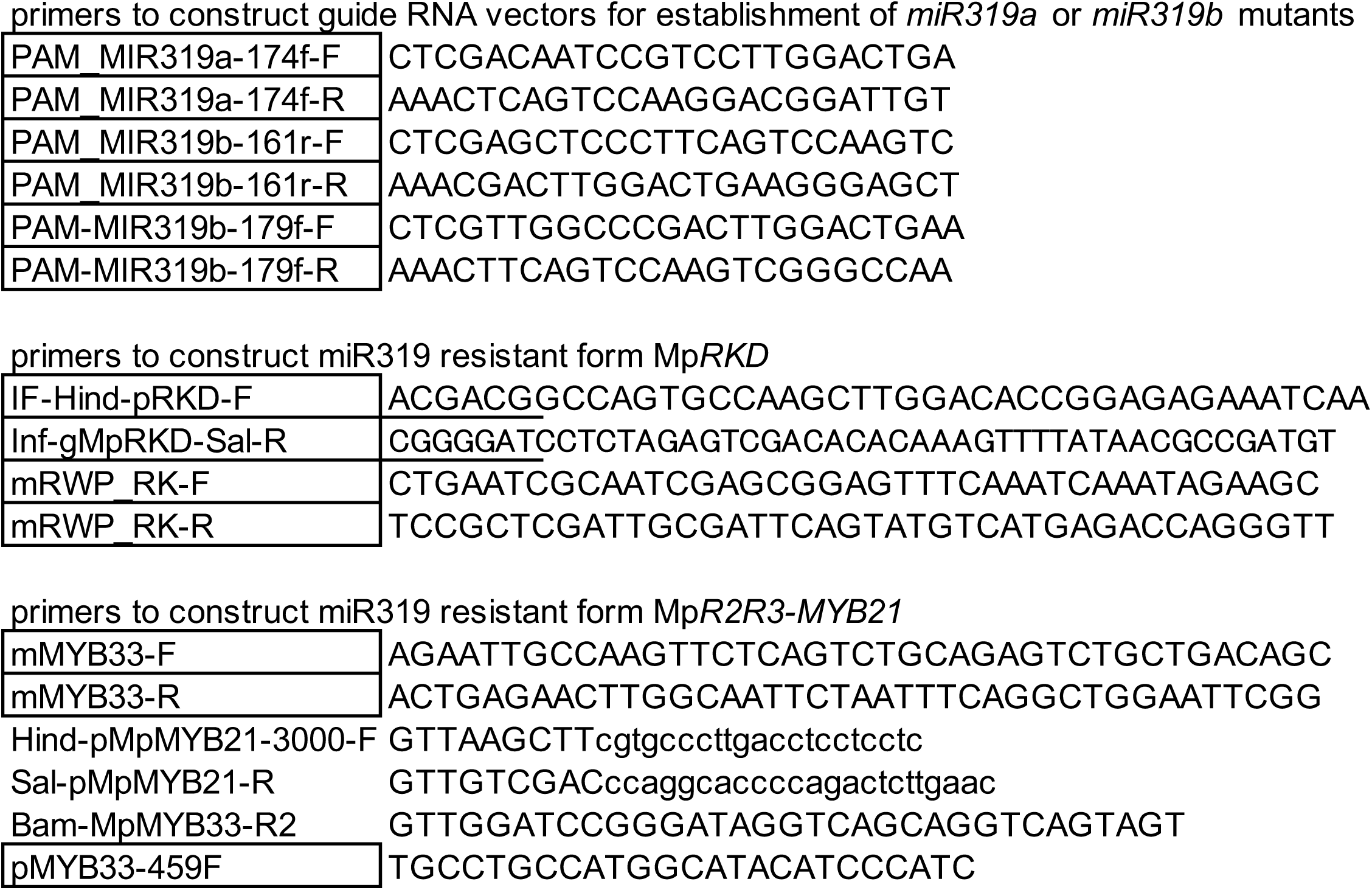
Primers used in this study.

**Fig.S1.**
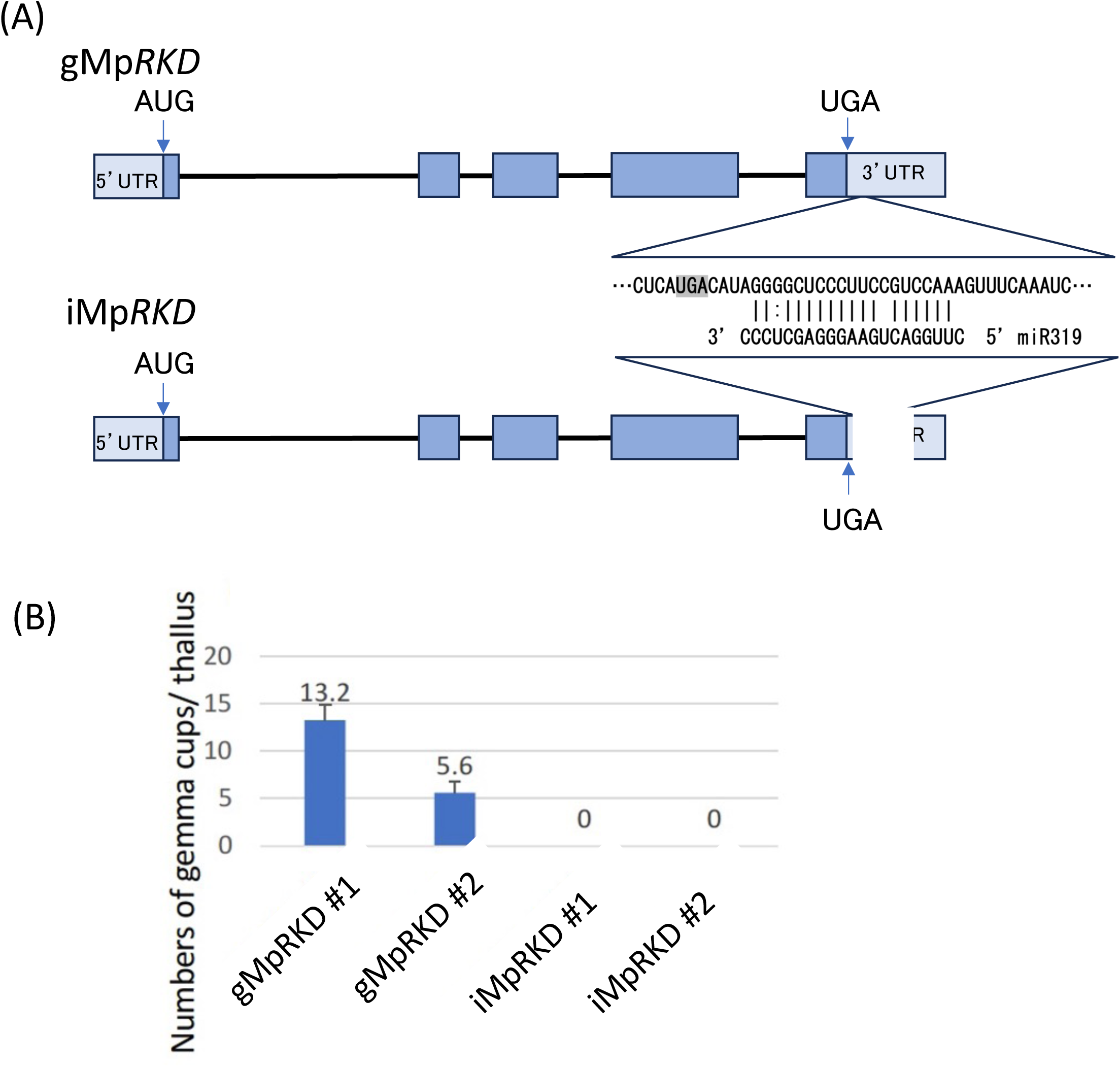
Introduction of the miR319-insensitive iMp*RKD* gene suppressed gemma cup formation in thalli. (A) Construct of wild-type *RKD* (gMp*RKD*) and miR319 insensitive *RKD* (iMp*RKD*) genes. The sequence segment complementary to miR319 sequences located in 3’ untranslated region (3’ UTR) was deleted to obtain the iMp*RKD* gene. Each construct was placed under the direction of the promoter of Mp*RKD* and introduced into Tak-1 wild type plants to obtain respective transformants. (B) The numbers of gemma cups were counted per thallus for transformants.

## Acknowledgements

We thank Kohchi, T. and Yamaoka, S. of Kyoto University, Ishizaki, K. of Kobe University, Shimamura, M. of Hiroshima University, and Nakajima, K. of NAIST for their advice and discussions. We also thank Bahari M. and Edanz (https://jp.edanz.com/ac) for editing a draft of this manuscript.

## Author contributions

KF performed the experiments. KF and MT designed the experiments. KF, YW, and MT wrote the manuscript.

## Data Availability

Relevant data can be found within the manuscript and its supplementary data online.

## References

Alaba S, Piszczalka P, Pietrykowska H, Pacak AM, Sierocka I, Nuc PW, Singh K, Plewka P, Sulkowska A, Jarmolowski A, et al (2015) The liverwort Pellia endiviifolia shares microtranscriptomic traits that are common to green algae and land plants. New Phytol 206: 352–367

Axtell MJ, Snyder JA, Bartel DP (2007) Common functions for diverse small RNAs of land plants. Plant Cell 19: 1750–1769

Bowman JLJL, Kohchi T, Yamato KT, Jenkins J, Shu S, Ishizaki K, Yamaoka S, Nishihama R, Nakamura Y, Berger F, et al (2017) Insights into Land Plant Evolution Garnered from the Marchantia polymorpha Genome. Cell 171: 287–304.e15

Gamborg OL, Miller RA, Ojima K (1968) Nutrient requirements of suspension cultures of soybean root cells. Exp Cell Res 50: 151–158

Hiwatashi T, Quan KL, Yasui Y, Takami H, Kajikawa M, Kirita H, Sato M, Wakazaki M, Yamaguchi K, Shigenobu S, et al (2019) The RopGEF KARAPPO Is Essential for the Initiation of Vegetative Reproduction in Marchantia polymorpha. Curr Biol 29: 3525–3531.e7

Ishizaki K, Chiyoda S, Yamato KT, Kohchi T (2008) Agrobacterium-mediated transformation of the haploid liverwort Marchantia polymorpha L., an emerging model for plant biology. Plant Cell Physiol 49: 1084–1091

Ishizaki K, Nishihama R, Ueda M, Inoue K, Ishida S, Nishimura Y, Shikanai T, Kohchi T (2015) Development of gateway binary vector series with four different selection markers for the liverwort marchantia polymorpha. PLoS One 10: 1–13

Koi S, Hisanaga T, Sato K, Shimamura M, Yamato KT, Ishizaki K, Kohchi T, Nakajima K (2016) An Evolutionarily Conserved Plant RKD Factor Controls Germ Cell Differentiation. Curr Biol 26: 1–7

Komatsu A, Kodama K, Mizuno Y, Fujibayashi M, Naramoto S, Kyozuka J (2023) Control of vegetative reproduction in Marchantia polymorpha by the KAI2-ligand signaling pathway. Curr Biol 33: 1196–1210.e4

Kosugi S, Ohashi Y (1997) PCF1 and PCF2 specifically bind to cis elements in the rice proliferating cell nuclear antigen gene. Plant Cell 9: 1607–1619

Koyama T, Sato F, Ohme-Takagi M (2017) Roles of miR319 and TCP transcription factors in leaf development. Plant Physiol 175: 874–885

Kubota A, Ishizaki K, Hosaka M, Kohchi T (2013) Efficient Agrobacterium-mediated transformation of the liverwort Marchantia polymorpha using regenerating thalli. Biosci Biotechnol Biochem 77: 167–172

Lin P-CC, Lu C-WW, Shen B-NN, Lee G-ZZ, Bowman JL, Arteaga-Vazquez MA, Liu L- YD, Hong S-FF, Lo C-FF, Su G-MM, et al (2016a) Identification of miRNAs and their targets in the liverwort Marchantia polymorpha by integrating RNA-Seq and degradome analyses. Plant Cell Physiol 57: 339–358

Lin P, Lu C, Shen B, Lee G, Bowman JL, Arteaga-vazquez MA, Liu LD, Hong S, Lo C, Su G, et al (2016b) Identification of miRNAs and Their Targets in the Liverwort Marchantia polymorpha by Integrating RNA-Seq and Degradome Analyses Special Focus Issue – Regular Paper. 57: 339–358

Lin SS, Bowman JL (2018) MicroRNAs in Marchantia polymorpha. New Phytol 220: 409–416

Martín-Trillo M, Cubas P (2010) TCP genes: a family snapshot ten years later. Trends Plant Sci 15: 31–39

Palatnik JF, Allen E, Wu X, Schommer C, Schwab R, Carrington JC, Weigel D (2003) Control of leaf morphogenesis by microRNAs. Nature 425: 257–263

Palatnik JF, Wollmann H, Schommer C, Schwab R, Boisbouvier J, Rodriguez R, Warthmann N, Allen E, Dezulian T, Huson D, et al (2007) Sequence and expression differences underlie functional specialization of Arabidopsis microRNAs miR159 and miR319. Dev Cell 13: 115–25

Pietrykowska H, Sierocka I, Zielezinski A, Alisha A, Carrasco-Sanchez JC, Jarmolowski A, Karlowski WM, Szweykowska-Kulinska Z (2022) Biogenesis, conservation, and function of miRNA in liverworts. J Exp Bot 73: 4528–4545

Rhoades MW, Reinhart BJ, Lim LP, Burge CB, Bartel B, Bartel DP (2002) Prediction of plant microRNA targets. Cell 110: 513–520

Romani F, Sauret-Güeto S, Rebmann M, Annese D, Bonter I, Tomaselli M, Dierschke T, Delmans M, Frangedakis E, Silvestri L, et al (2024) The landscape of transcription factor promoter activity during vegetative development in Marchantia. Plant Cell 36: 2140–2159

Rövekamp M, Bowman JL, Grossniklaus U (2016) Marchantia MpRKD Regulates the Gametophyte-Sporophyte Transition by Keeping Egg Cells Quiescent in the Absence of Fertilization. Curr Biol 26: 1782–1789

Shimamura M (2016) Marchantia polymorpha: Taxonomy, phylogeny and morphology of a model system. Plant Cell Physiol 57: 230–256

Solly JE, Cunniffe NJ, Harrison CJ (2017) Regional Growth Rate Differences Specified by Apical Notch Activities Regulate Liverwort Thallus Shape. Curr Biol 27: 16–26

Sugano SS, Nishihama R, Shirakawa M, Takagi J, Matsuda Y, Ishida S, Shimada T, Hara-Nishimura I, Osakabe K, Kohchi T (2018) Efficient CRISPR/Cas9-based genome editing and its application to conditional genetic analysis in Marchantia polymorpha. PLoS One 13: e0205117

Sugano SS, Shirakawa M, Takagi J, Matsuda Y, Shimada T, Hara-Nishimura I, Kohchi T (2014) CRISPR/Cas9-mediated targeted mutagenesis in the liverwort Marchantia polymorpha L. Plant Cell Physiol 55: 475–481

Tsuboyama K, Tadakuma H, Tomari Y (2018a) Conformational Activation of Argonaute by Distinct yet Coordinated Actions of the Hsp70 and Hsp90 Chaperone Systems. Mol Cell 70: 722–729.e4

Tsuboyama S, Kodama Y (2018) Highly efficient G-AgarTrap-mediated transformation of the Marchantia polymorpha model strains Tak-1 and Tak-2. Plant Biotechnol 35: 399–403

Tsuboyama S, Nonaka S, Ezura H, Kodama Y (2018b) Improved G-AgarTrap: A highly efficient transformation method for intact gemmalings of the liverwort Marchantia polymorpha. Sci Rep 8: 1–10

Tsuzuki M, Futagami K, Shimamura M, Inoue C, Kunimoto K, Oogami T (2019) An Early Arising Role of the MicroRNA156 / 529-SPL Module in Reproductive Development Revealed by the Liverwort Marchantia polymorpha Report An Early Arising Role of the MicroRNA156 / 529-SPL Module in Reproductive Development Revealed by the Liverwort. Curr Biol 29: 1–8

Tsuzuki M, Nishihama R, Ishizaki K, Kurihara Y, Matsui M, Bowman JLJL, Kohchi T, Hamada T, Watanabe Y (2016) Profiling and characterization of small RNAs in the liverwort, Marchantia polymorpha, belonging to the first diverged land plants. Plant Cell Physiol 57: 359–372

Yamaoka S, Inoue K, Araki T (2021) Regulation of gametangia and gametangiophore initiation in the liverwort Marchantia polymorpha. Plant Reprod. doi: 10.1007/s00497-021-00419-y

Yasui Y, Tsukamoto S, Sugaya T, Nishihama R, Wang Q, Kato H, Yamato KT, Fukaki H, Mimura T, Kubo H, et al (2019) GEMMA CUP-ASSOCIATED MYB1, an Ortholog of Axillary Meristem Regulators, Is Essential in Vegetative Reproduction in Marchantia polymorpha. Curr Biol 29: 3987–3995.e5

